# An integrated design strategy for developing and validating microalgal formulations in common bean and rainfed rice

**DOI:** 10.64898/2026.07.27.740880

**Authors:** Carlos Lopera, Madelem Giraldo, Natalia Herrera

## Abstract

Microalgae and cyanobacteria have emerged as promising resources for sustainable agriculture; however, integrated methodologies for the rational design of crop-specific agricultural formulations remain scarce. This study proposes an integrated framework that combines biomass production, species characterization, nutrient characterization, mixture design, nutrient profile estimation, biological validation, and statistical optimization for the rational development of agricultural formulations based on microalgae and cyanobacteria. As a proof of concept, the proposed framework was applied to formulate consortia composed of *C*. *vulgaris*, *Scenedesmus* sp., and *A*. *platensis* using common bean (*Phaseolus vulgaris* L., ecotype ‘Sangre Toro’) and rainfed rice (*Oryza sativa* L., cv. ‘Fedearroz 2020’) as model crops. The experimentally determined nutrient composition of the individual biomasses was integrated into a simplex-lattice mixture design coupled with response surface methodology and desirability analysis to identify crop-specific optimal formulations and estimate their nutrient profiles. The cubic model provided the best fit (P < 0.05), showing high predictive performance and a non-significant lack of fit. The optimal bean formulation consisted of 31.6% *C*. *vulgaris* and 68.4% *Scenedesmus* sp., whereas the optimal rice formulation comprised 62.3% *A*. *platensis* and 37.7% *C*. *vulgaris*, demonstrating distinct crop-specific responses. The optimized bean formulation exhibited higher estimated concentrations of calcium, phosphorus, iron, and zinc, whereas the rice formulation showed higher estimated potassium and Kjeldahl nitrogen contents. These findings demonstrate the feasibility of developing crop-specific microalgal formulations and highlight that different crops may require distinct formulations rather than a universal approach. The proposed approach offers a reproducible, integrated framework for the development and optimization of next-generation agricultural formulations for sustainable crop production.

## 1. INTRODUCTION

The rapid increase in the global population has intensified food demand, requiring more efficient and sustainable crop management strategies. To maximize productivity, agriculture has become highly dependent on inorganic fertilizers; however, their excessive use has contributed to soil degradation, water contamination, greenhouse gas emissions, and the loss of long-term agricultural sustainability [1,2]. In addition, inadequate soil management practices have reduced soil biological activity and nutrient cycling, limiting the capacity of agricultural systems to meet increasing food demands [3]. Among the alternatives proposed for sustainable agriculture, microalgae and cyanobacteria have emerged as promising biological resources for developing biofertilizers and biostimulants [4].

These microorganisms synthesize phytohormones, amino acids, vitamins, polysaccharides, and other bioactive compounds that stimulate plant growth while also supplying macro- and micronutrients [2,3]. Their application has been associated with improved nutrient availability, seed germination, plant physiological performance, tolerance to abiotic stress, and beneficial changes in soil structure and microbial communities, supporting circular bioeconomy strategies for agriculture [5–7].

Despite these advances, the development of microalgal formulations remains largely empirical [3,8]. Most studies focus on evaluating individual strains or predefined consortia without integrating biomass characterization, formulation design, nutrient composition, statistical optimization, and biological validation into a unified development strategy. Consequently, reproducible methodologies for the rational design of crop-specific agricultural formulations remain scarce.

To address this gap, this study proposes an integrated approach for the rational design of agricultural formulations based on microalgae and cyanobacteria. The framework combines biomass production, species characterization, nutritional characterization, mixture design, nutrient profile estimation, biological validation, and statistical optimization into a single framework for formulation development and optimization.

As a proof of concept, formulations composed of *C*. *vulgaris*, *Scenedesmus* sp., and *A*. *platensis* were designed using a simplex–lattice mixture design coupled with response surface methodology. Common bean (*Phaseolus vulgaris* L., ecotype ‘Sangre Toro’) and rainfed rice (*Oryza sativa* L., cv. ‘Fedearroz 2020’) were selected as model crops representing legumes and cereals, respectively. The nutritional characterization of each biomass was integrated into the formulation process to estimate the nutrient composition of the optimized mixtures, while biological validation was performed by evaluating crop biometric responses.

Unlike conventional workflows, the proposed design strategy uses biomass characterization as quantitative input for formulation design and statistical optimization, providing a reproducible, integrated design strategy for developing crop-specific formulations. The proposed framework is summarized in Figure 1.

**Fig. 1.**
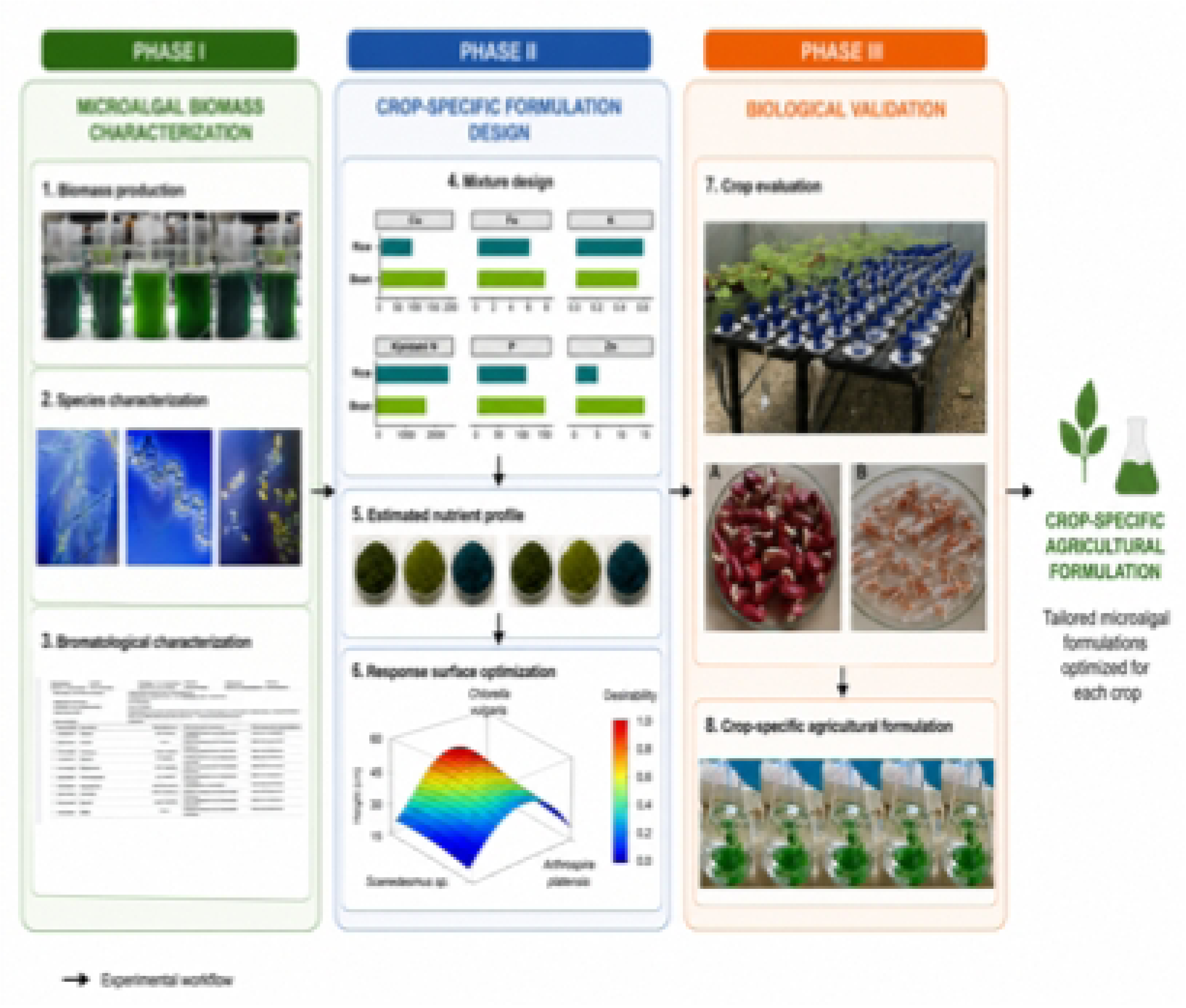
Integrated framework for the rational design of crop-specific agricultural formulations based on microalgae and cyanobacteria. The design strategy is organized into three phases: (I) microalgal biomass characterization, including biomass production, species identification, and nutritional characterization; (II) crop-specific formulation design through mixture design, nutrient profile estimation, and statistical optimization; and (III) biological validation leading to the selection of optimal agricultural formulations.

**Fig. 2.**
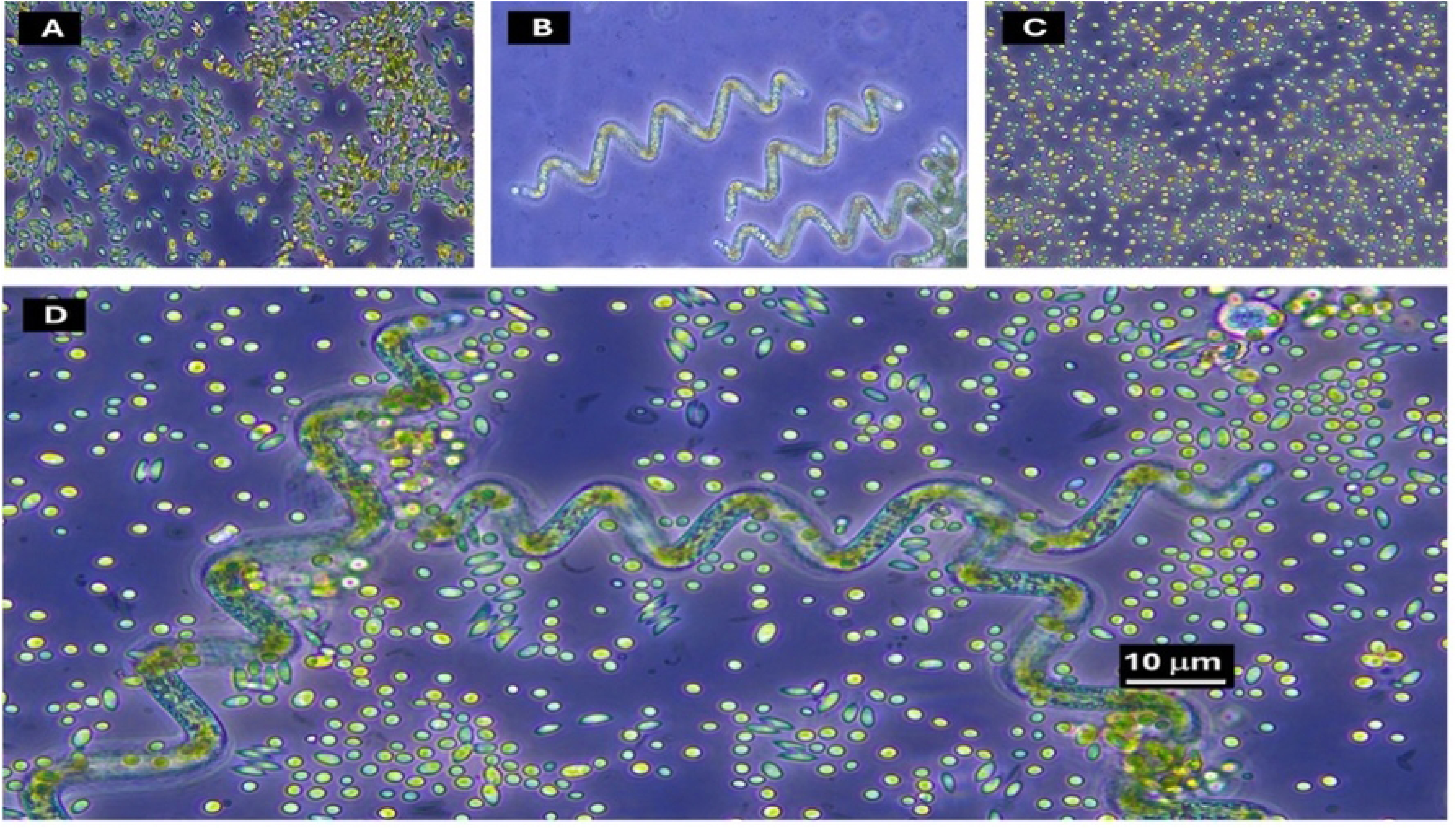
Inverted phase-contrast microscope photographs (40X) of the microalgae applied to the crops. A. *Scenedesmus* sp*; B. A. platensis*; C. *C*. *vulgaris,* and D. Mixture with the three species of microalgae.

## 2. METHODOLOGY

### 2.1. Microalgae and cultivation

The biomass of the microalgae *C. vulgaris*, *Scenedesmus* sp., and *A. platensis* was supplied in liquid suspension by the Natural Products Laboratory of the University of Antioquia. *C. vulgaris* and *Scenedesmus* sp. were cultivated in BBM medium (pH 7.5) and *A. platensis* in Zarrouk medium (pH 9.5). The cultures were initially carried out in 500 mL Erlenmeyer flasks fitted with a sterilizable rubber stopper with two outlets: one to maintain constant aeration at 1 L min⁻¹ and the other to release pressure from the culture. This system was kept on open shelves at 25 ± 1 °C, under a 12-hour light/12-hour darkness photoperiod. Cultures were renewed every 12 days by adding 10 mL inoculum to 400 mL fresh medium [9]. Subsequently, the culture was scaled up to 20 L in closed cylindrical bioreactors. Culture conditions were as follows: a 12 h light/12 h dark photoperiod, a temperature of 25 ± 1 °C, a light intensity of approximately 268–376 µmol photons m⁻² s⁻¹, estimated from 18,790 lux using published conversion factors for white LED lighting, a pH of 7.0 ± 0.5 for BBM media, and a pH of 9.0 ± 0.5 for Zarrouk medium. Cultures were continuously aerated and mixed with filtered air supplied by the centralized medical air system of the University Research Center under standardized laboratory conditions. All cultures were maintained for 20 days prior to biomass harvesting and subsequent nutritional characterization.

### 2.2. Microalgae harvest

*A. platensis* was harvested after 20 days by filtration with a 100 mm mesh. In the case of *C*. *vulgaris* and *Scenedesmus* sp., biomass separation was achieved by sedimentation, eliminating the need for aeration after 20 days, allowing the biomass to accumulate at the bottom of the bioreactor. Finally, cell concentration was estimated by optical density (OD 630). Optical density measurements were correlated with dry biomass using calibration curves established for each microalgal species. 300 mL of culture was taken, transferred to 96-well plates, and measured at 630 nm with a Multiskan Spectrum spectrophotometer (Thermo Scientific). Measurements were performed in triplicate.

### 2.3. Nutritional characterization of microalgal and cyanobacterial biomass

Microalgal biomass was produced in 3-L Erlenmeyer flasks containing 2.5 L of culture medium, using 400 mL of a 12-day-old inoculum for each species. Cultures were maintained under the culture conditions described in the previous section for 20 days, with continuous aeration at 1 L min⁻¹. Biomass was harvested by sedimentation for *C. vulgaris* and *Scenedesmus* sp., whereas *A*. *platensis* was harvested by filtration through a 15-µm mesh. The collected biomass was washed with Milli-Q water, freeze-dried (LABCONCO FreeZone 12L), and dry weight was determined. The freeze-dried biomass of each species was analyzed to determine its nutritional composition, including total oxidizable organic carbon (NTC 5167), Kjeldahl nitrogen (ISO 5983), phosphorus (NTC 4981), and the mineral elements calcium, copper, iron, magnesium, manganese, potassium, sodium, and zinc by atomic absorption spectrophotometry (NTC 5151). The quantified mineral elements correspond to nutrients accumulated by the biomass during cultivation following their uptake from the culture medium. The resulting nutritional data were subsequently used to estimate the nutrient composition of the optimized agricultural formulations. The nutritional composition of the optimal formulations was estimated from the experimentally determined nutrient composition of each microalgal species and the optimized species proportions obtained from the mixture design.

For each nutrient, the expected concentration in the formulation was calculated as the weighted average of the contribution of each species according to its proportion in the consortium. The estimated nutrient profiles of the optimal formulations for common bean and rice were visualized in R (version 2025.05.0+496) using the ggplot2 package.

### 2.4. Experimental Design and Optimization of Agricultural Formulations

The experiment was conducted in greenhouse facilities at Ecosphaira (Envigado, Antioquia, Colombia), located at 1,750 m above sea level in the rural area of the El Salado neighborhood.

To determine the optimal proportion of liquid microalgae biomass, STATGRAPHICS Centurion XIX statistical software was used to implement a simplex–lattice mixture design augmented with three components (independent variables): *C*. *vulgaris* (A), *Scenedesmus* sp. (B), and *A*. *platensis* (C). There were no restrictions on the inclusion levels of the mixture components, which were expressed as fractions of the mixture (A + B + C = 1). A total of 14 mixtures (formulations) were generated (Table 1). To measure the biometric variables (response variables), two indicator plant species were used: the bean (*Phaseolus vulgaris*) ecotype “Sangre toro” and rainfed rice variety Fedearroz 2020 (*Oryza sativa*). Each mixture was evaluated using independent experimental units, with a total of 28 replicates per crop species distributed across the 14 formulations, resulting in 56 experimental units in total.

**Table 1.** Simplex–lattice mixture design used for the development of agricultural formulations based on *C*. *vulgaris*, *Scenedesmus* sp., and *A. platensis*.

| *RUN* | *BLOCK* | *Components* |  |  |
| --- | --- | --- | --- | --- |
|  |  | *Chlorella* | *Scenedesmus* sp. | *Arthrospira platensis* |
|  |  | *vulgaris* |  |  |
| 1 | 1 | 0.666667 | 0.000000 | **0.333333** |
| 2 | 1 | 0.666667 | 0.166667 | **0.166667** |
| 3 | 1 | 0.000000 | 0.333333 | **0.666667** |
| 4 | 1 | 0.000000 | 1.000000 | **0.000000** |
| 5 | 1 | 0.0000 | 0.666667 | **0.333333** |
| 6 | 1 | 0.333333 | 0.000000 | **0.666667** |
| 7 | 1 | 0.333333 | 0.333333 | **0.333333** |
| 8 | 1 | 0.000000 | 0.000000 | **1.000000** |
| 9 | 1 | 1.000000 | 0.000000 | **0.000000** |
| 10 | 1 | 0.166667 | 0.166667 | **0.666667** |
| 11 | 1 | 0.166667 | 0.666667 | **0.166667** |
| 12 | 1 | 0.333333 | 0.666667 | **0.000000** |
| 13 | 1 | 0.333333 | 0.333333 | **0.333333** |
| 14 | 1 | **0.666667** | **0.333333** | **0.000000** |

### 2.5. Substrate and plant material

The substrate used to grow indicator plants in this experiment was obtained from a local commercial supplier and consisted of a 70%–30% (w/w) mixture of AFM-1 Germination Mix and medium-coarse vermiculite, providing a defined supply of macro- and micronutrients, including nitrogen (as NO₃⁻ and NH₄⁺), phosphorus, potassium, calcium, magnesium, and trace elements such as Zn and Mn, along with a mineral matrix rich in silicates (SiO₂, Al₂O₃, MgO) that supports structural stability and nutrient retention.

Seeds of each plant species were purchased from Asofril and Fedearroz. The seeds were pre- germinated in a humid chamber in an incubator (Memmert, Germany) at 30 °C for 4 days. Subsequently, two seeds per pot were sown in 600 mL plastic pots, and 10 days later, seedlings were thinned to one per pot by selecting the best-developed plant. The plants were grown under controlled conditions in a greenhouse (relative humidity: 82-87%; temperature: 22-25°C), and the substrate moisture was maintained between 50% and 60% of its maximum moisture retention capacity. They were fed weekly with Hoagland’s nutrient solution [10]

### 2.6. Preparation and application of mixtures

The liquid suspension biomass of each microalga was used to formulate the different mixtures (Table 1). Each pot received two applications of 5 mL of the corresponding mixture. This contained: *A. platensis*: 1 × ^cells/mL,^ *Scenedesmus* sp.: 3.3 ×^10⁸^ cells/mL, and *C. vulgaris*: 8.3 ×^10⁸^ cells/mL. The first application was made 10 days after sowing, and the second 15 days after the first application. The evaluated microalgal formulations, composed of *Scenedesmus* sp., *A*. *platensis*, and *C*. *vulgaris*, were examined using an inverted phase-contrast microscope (Leica). Conventional wet mounts were prepared by placing an aliquot of each sample on a glass microscope slide, covered with a coverslip, and observed under phase-contrast illumination at magnifications of up to 40× to assess the morphology and distribution of the microalgae and cyanobacteria.

### 2.7. Measurement of biometric variables

#### 2.7.1 “Sangre toro” bean

When the plants reached the pre-flowering stage, 40 days after sowing, at the time when the flower bud was observed, the following biometric variables were determined: (i) plant height, measured as the distance between the base of the stem and the last meristem; (ii) leaf diameter, measured longitudinally on the terminal leaflet of the third true (compound) leaf using a millimeter ruler; (iii) stem diameter, measured at the middle third of the stem with a caliper (MP Tools); and (iv) total dry biomass (TDB): the aerial and root parts of each plant were harvested, transferred to paper bags, and dried at 72°C in a dehydrator (Innova) until a constant weight was obtained. To harvest the aerial part, the stem was cut flush with the substrate. To harvest the root, the substrate was removed from the pot and washed several times with distilled water to remove excess substrate. The sum of the aerial biomass and root biomass was recorded as total dry biomass.

#### 2.7.2. Fedearroz 2020 rainfed rice variety

When the plants reached the final primordium stage, 60 days after sowing, the following biometric variable was determined: total dry biomass (TDB). As described for bean plants, the aerial and root tissues were harvested, transferred to paper bags, and dehydrated at 72°C until a constant weight was obtained. The sum of the above-ground biomass and root biomass was recorded as total dry biomass.

### 2.8. Validation of the mixture

To validate the optimal microalgae mixture and its biostimulant effects, a second experiment was conducted in a greenhouse using a completely randomized design with a single-factor arrangement of treatments. The factor evaluated was the microalgae dose with two levels or treatments: i) (+) microalgae, the plants received 5 mL of the optimal microalgae mixture; ii) control, the plants did not receive microalgae. The treatments were applied to the same indicator plants used for the mixture design, and the same procedures were followed. Ten replicates were used for each plant species. Before performing the analysis of variance (ANOVA) to test the hypothesis that microalgae exert biostimulant effects on “Sangre toro” bean and rice plants, the assumptions of normality of residuals and homogeneity of variances were evaluated. Normality and homoscedasticity were assessed using the Shapiro Wilk W test and Levene’s test, respectively.

### 2.9. Data analysis

Parameters such as sum of squares, mean square, degrees of freedom, F values, p values, and coefficients of determination (R2) were obtained from an analysis of variance (ANOVA) generated by the statistical software STATGRAPHICS Centurion XIX. The statistical significance of the models was determined at the 5% significance level (α = 0.05), assessed using the F-ratio. The adjusted (R2-adj) and predicted (R2-pred) coefficients of determination were used to compare the different models and select an appropriate model for each response. Response surface graphs generated from the selected models were used to study the interactive effects of the components (microalgae) on the responses. The objective of this statistical approach was to identify the microalgae mixture that maximized the biometric variables studied. The optimal mixture was validated, and biostimulant effects were investigated using a one-way analysis of variance in STATGRAPHICS Centurion XIX.

## 3. RESULTS

### 3.1. Microalgal culture and species characterization

Each 20 L photobioreactor produced approximately 18 g of dry biomass over a 20-day cultivation period. Microscopic examination confirmed the presence of the three taxa included in the formulations according to their diagnostic morphological features. *Scenedesmus* sp. exhibited elongated cells with pointed apical poles, characteristic of the genus. *Arthrospira platensis* displayed characteristic helically coiled trichomes composed of cylindrical cells; and *C*. *vulgaris* appeared as unicellular spherical to subspherical coccoid cells with smooth cell walls. The simultaneous observation of these morphological traits confirmed the taxonomic identity of the species composing the consortium [11,12].

### 3.2. Nutrient composition of microalgal biomass

The nutrient composition of the evaluated biomasses revealed distinct elemental profiles among the three species (Fig. 3). *C*. *vulgaris* and *Scenedesmus* sp. exhibited the highest calcium concentrations (190 and 197 mg kg⁻¹, respectively), whereas *A*. *platensis* contained only 19 mg kg⁻¹. Likewise, phosphorus concentrations were higher in *Scenedesmus* sp. (171 mg kg⁻¹) and *C*. *vulgaris* (142 mg kg⁻¹), while *A*. *platensis* contained <100 mg kg⁻¹. Iron concentrations reached 8 mg kg⁻¹ in both *C*. *vulgaris* and *Scenedesmus* sp., whereas *A*. *platensis* contained <5 mg kg⁻¹. Zinc was detected only in *C*. *vulgaris* and *Scenedesmus* sp. (15 mg kg⁻¹), while it was not detected in *A*. *platensis*. Copper was below the detection limit (<5 mg kg⁻¹) in both green microalgae and was not detected in *A*. *platensis*. Magnesium and manganese remained below 50 mg kg⁻¹ and 5 mg kg⁻¹, respectively, in all evaluated biomasses. Sodium concentrations were also below 500 mg kg⁻¹ in the three species.

**Fig. 3.**
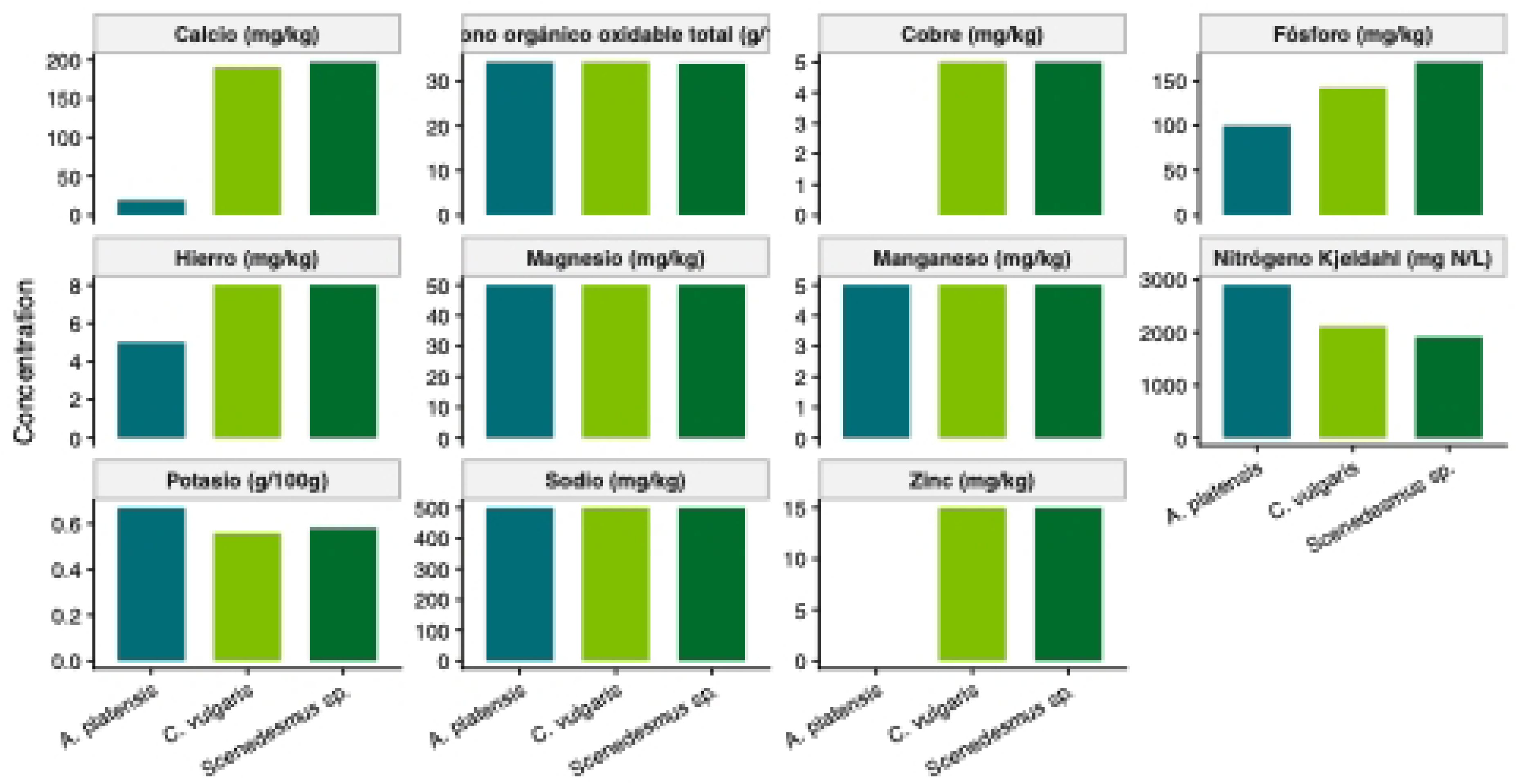
Nutrient composition of the biomass of *A*. *platensis*, *C*. *vulgaris*, and *Scenedesmus* sp., showing the concentrations of macro and micronutrients used for estimating the nutrient composition of the optimized agricultural formulations. Bars represent the concentrations of calcium (Ca), phosphorus (P), iron (Fe), zinc (Zn), potassium (K), and Kjeldahl nitrogen in each formulation, calculated from the experimentally determined nutritional composition of the individual microalgal species and their respective proportions in each consortium. Calcium, phosphorus, iron, and zinc are expressed as mg kg⁻¹ dry biomass; potassium as g 100 g⁻¹ dry biomass; and Kjeldahl nitrogen as mg N L⁻¹.

In contrast, *A*. *platensis* exhibited the highest potassium concentration (0.67 g 100 g⁻¹) and the highest Kjeldahl nitrogen content (2897 mg N L⁻¹), followed by *C*. *vulgaris* (2104 mg N L⁻¹) and *Scenedesmus* sp. (1913 mg N L⁻¹). Total oxidizable organic carbon showed very similar values among species, ranging from 34.0 to 34.3 g 100 g⁻¹.

### 3.3. Crop-specific optimal formulations and estimated nutrient profiles

The simplex–lattice mixture design identified different optimal formulations for common bean and rainfed rice, confirming that the optimal biomass composition depended on the crop species (Fig. 4). The formulation predicted for common bean consisted of 68.4% *Scenedesmus* sp. and 31.6% *C*. *vulgaris*, whereas *A*. *platensis* was not included in the optimized mixture. In contrast, the optimal formulation for rainfed rice comprised 62.3% *A*. *platensis* and 37.7% *C*. *vulgaris*, with no contribution from *Scenedesmus* sp.

**Fig. 4.**
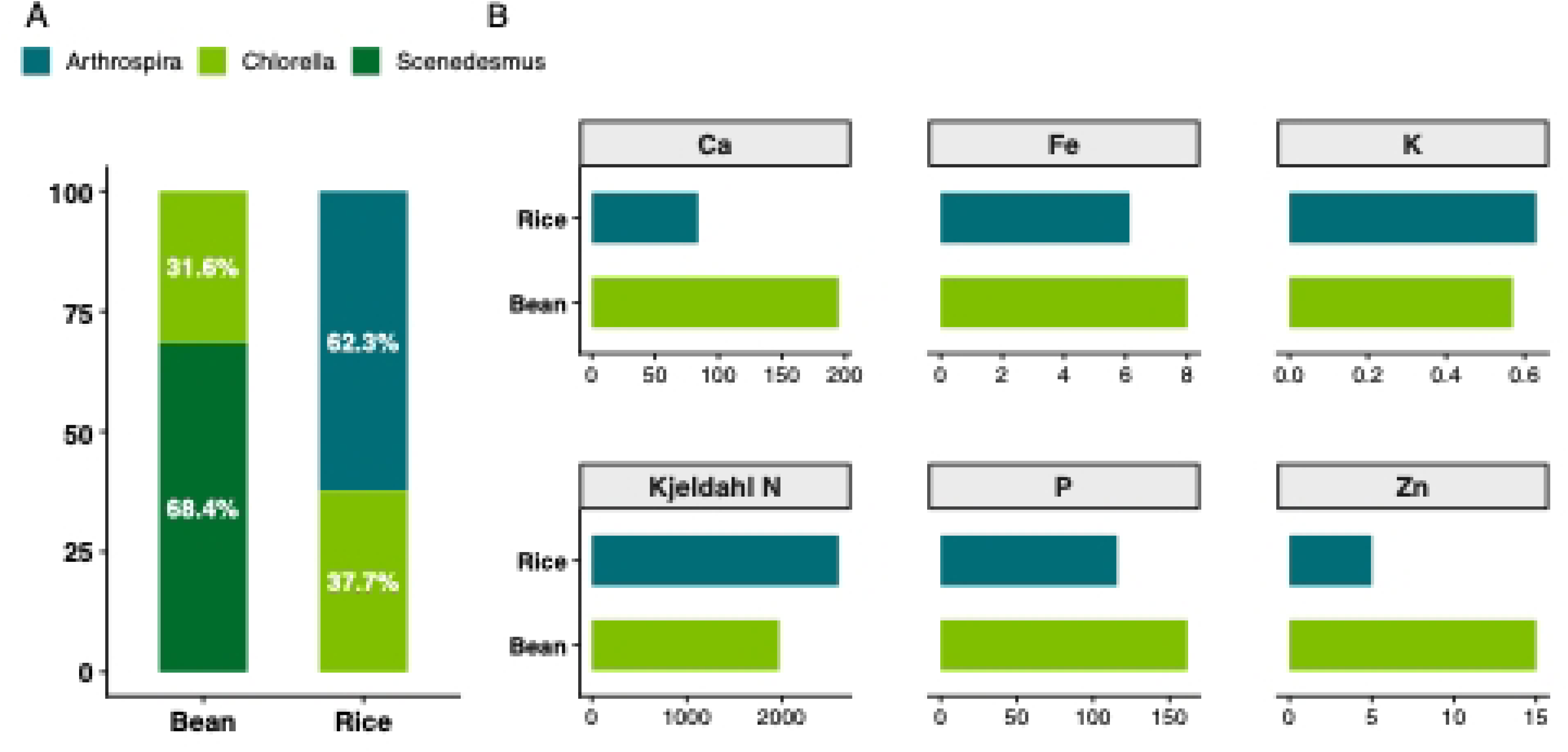
Predicted optimal agricultural formulations for bean and rainfed rice and the estimated concentrations of selected nutrients in each formulation. (A) Optimal species composition. (B) Estimated nutrient profiles.

These differences in species composition resulted in distinct estimated nutrient profiles between formulations. The formulation optimized for common bean presented higher estimated concentrations of calcium (190 mg kg⁻¹), iron (8 mg kg⁻¹), phosphorus (approximately 150 mg kg⁻¹), and zinc (15 mg kg⁻¹) than the formulation optimized for rice. Conversely, the rice formulation exhibited higher estimated concentrations of potassium (0.67 g 100 g⁻¹) and Kjeldahl nitrogen (approximately 2,300 mg N L⁻¹), reflecting the greater contribution of *A*. *platensis*.

### 3.4. “Sangre toro” bean

#### 3.4.1. Model performance and optimization

The cubic model provided the best fit for all evaluated variables (P < 0.05), explaining 89.1%, 90.7%, 87.4%, and 75.2% of the observed variability in plant height, total dry biomass (TDB), leaf diameter, and stem diameter, respectively (S1 File. Table S1). Significant relationships were found between the proportions of *C*. *vulgaris*, *Scenedesmus* sp., and *A*. *platensis* and all biometric variables (S1 File. Table S2). Response surface analysis showed that the formulations maximizing plant height, TDB, and stem diameter were characterized by a higher proportion of *Scenedesmus* sp. and the absence of *A*. *platensis*, whereas leaf diameter was maximized by a formulation enriched in *A*. *platensis* without *C*. *vulgaris* (Table 2; Fig. 5).

**Fig. 5.**
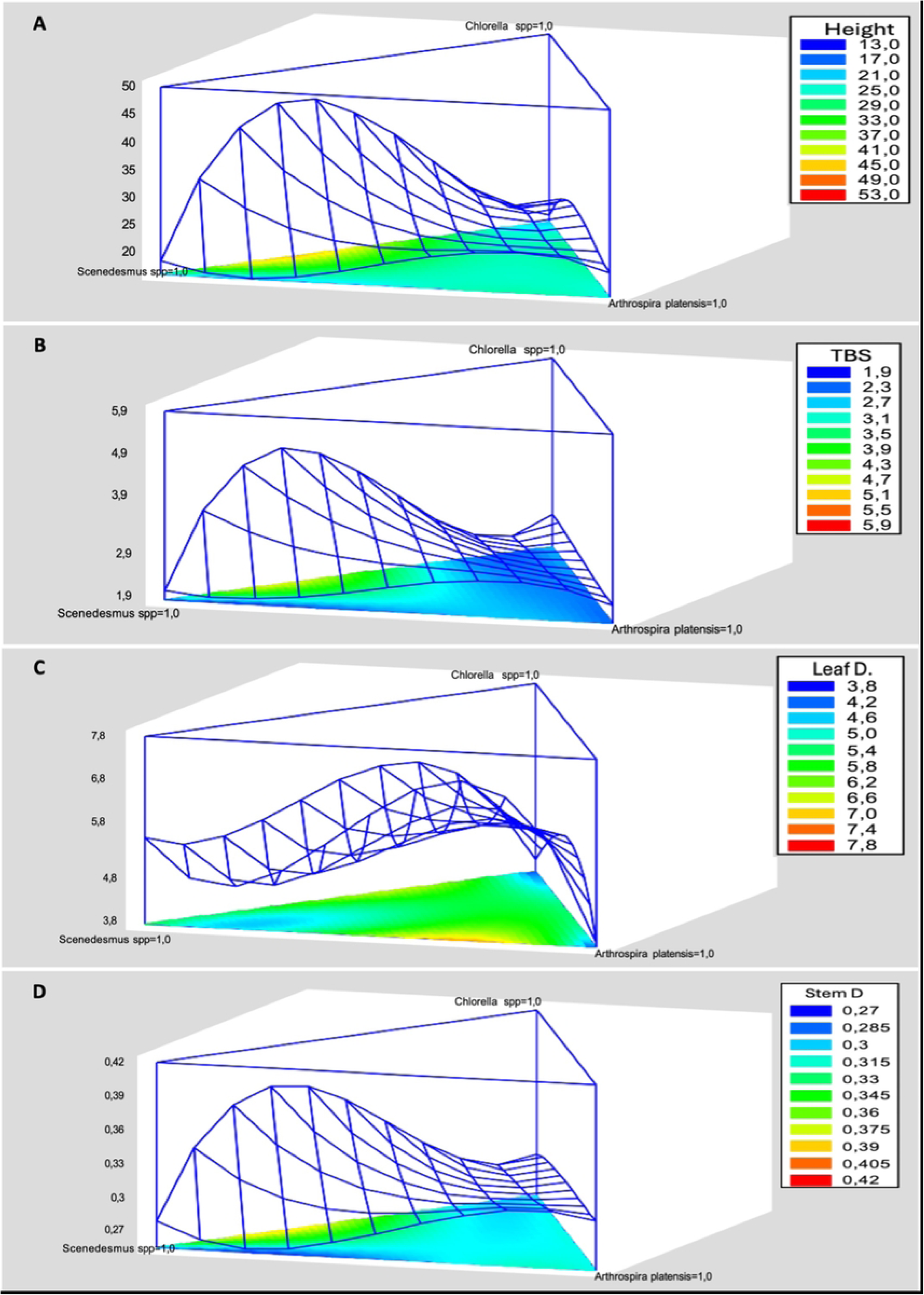
Response surface plots illustrating the effects of microalgal mixture composition (*C*. *vulgaris*, *Scenedesmus* sp., and *A*. *platensis*) on the evaluated variables in “Sangre Toro” bean plants. A: plant height; B: total dry biomass (TDB); C: leaf diameter; D: stem diameter.

**Table 2.** Predicted optimal agricultural formulations obtained from the simplex–lattice mixture design, showing the component proportions that maximized each biometric response variable.

| *Responses* | *Optimum* | *Chlorella* | *Scenedesmus* sp. | *Arthrospira platensis* |
| --- | --- | --- | --- | --- |
|  | *value* | *vulgaris* |  |  |
| **Height** | 45.0578 | 0.338713 | 0.661287 | **9.77515E-11** |
| **TDB** | 4.76111 | 0.298251 | 0.701749 | **2.02883E-10** |
| **Leaf diameter** | 7.19041 | 2.49336E-10 | 0.298951 | **0.701049** |
| **Stem diameter** | **0.387765** | **0.315072** | **0.684928** | **1.45122E-9** |

#### 3.4.2. Multiple response optimization and biological validation

Simultaneous optimization using the desirability function identified a single formulation that maximized the overall biometric performance. The optimal formulation consisted of 31.6% *C*. *vulgaris*, 68.4% *Scenedesmus* sp., with no contribution from *A*. *platensis*, reaching an overall desirability value of 0.89 (Fig. 6). The model predicted values of 45.03 cm for plant height, 4.75 g plant⁻¹ for TDB, 6.74 cm for leaf diameter, and 0.38 cm for stem diameter.

**Fig. 6.**
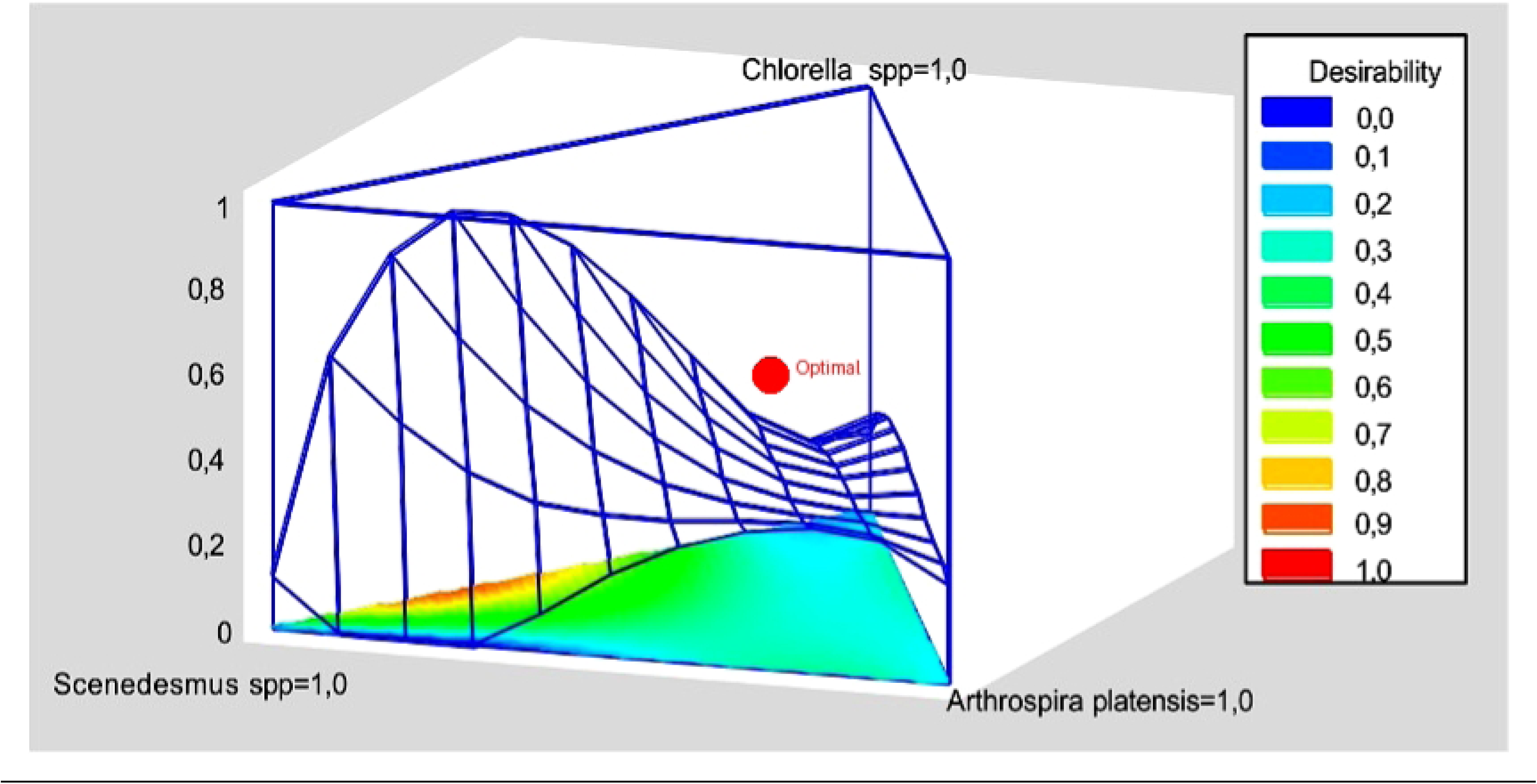
Desirability response surface showing the optimal microalgal mixture and predicted responses for the evaluated variables.

Experimental verification confirmed the predictive performance of the model, as experimental values (46.42 cm plant⁻¹, 4.80 g plant⁻¹, 6.81 cm, and 0.40 cm, respectively) closely matched the predicted responses. Residual normality and homogeneity of variances were verified for all variables (P ≥ 0.05), with a Box–Cox transformation applied only to plant height.

Application of the optimized formulation produced a significant biostimulant effect (P < 0.05), increasing plant height, TDB, leaf diameter, and stem diameter by 14.8%, 13.1%, 12.5%, and 11.8%, respectively, compared with the control (Fig. 7). These improvements were also visually evident through greater shoot growth and root biomass development in treated plants (Fig. 8).

**Fig. 7.**
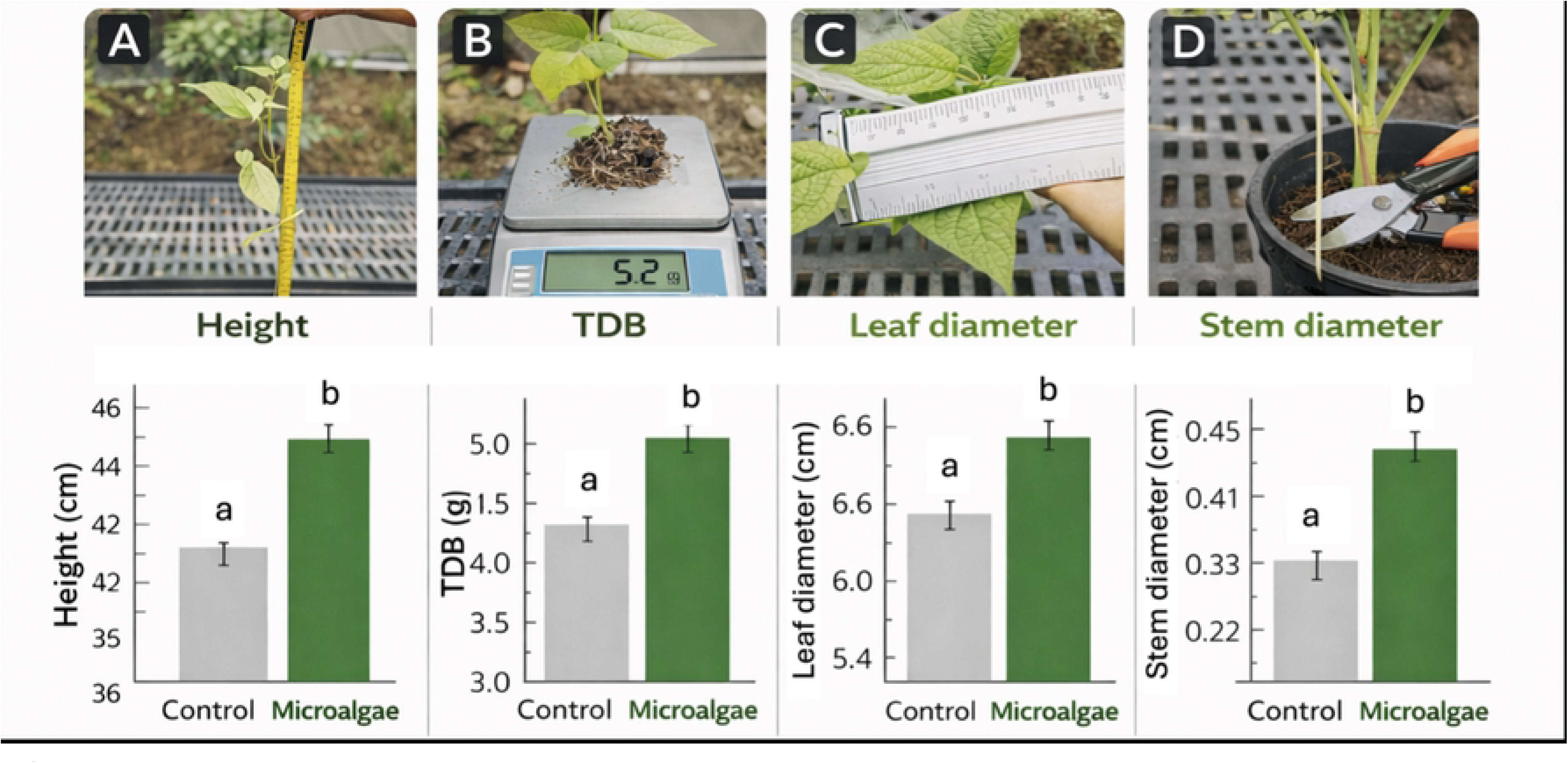
Biostimulant effect of microalgae on biometric variables in “Sangre Toro” bean plants. A: height; B: TDB; C: leaf diameter; D: stem diameter. The vertical bars indicate the standard error of the mean (n=10). Averages with different letters differ significantly according to ANOVA (P ≤ 0.05).

**Fig. 8.**
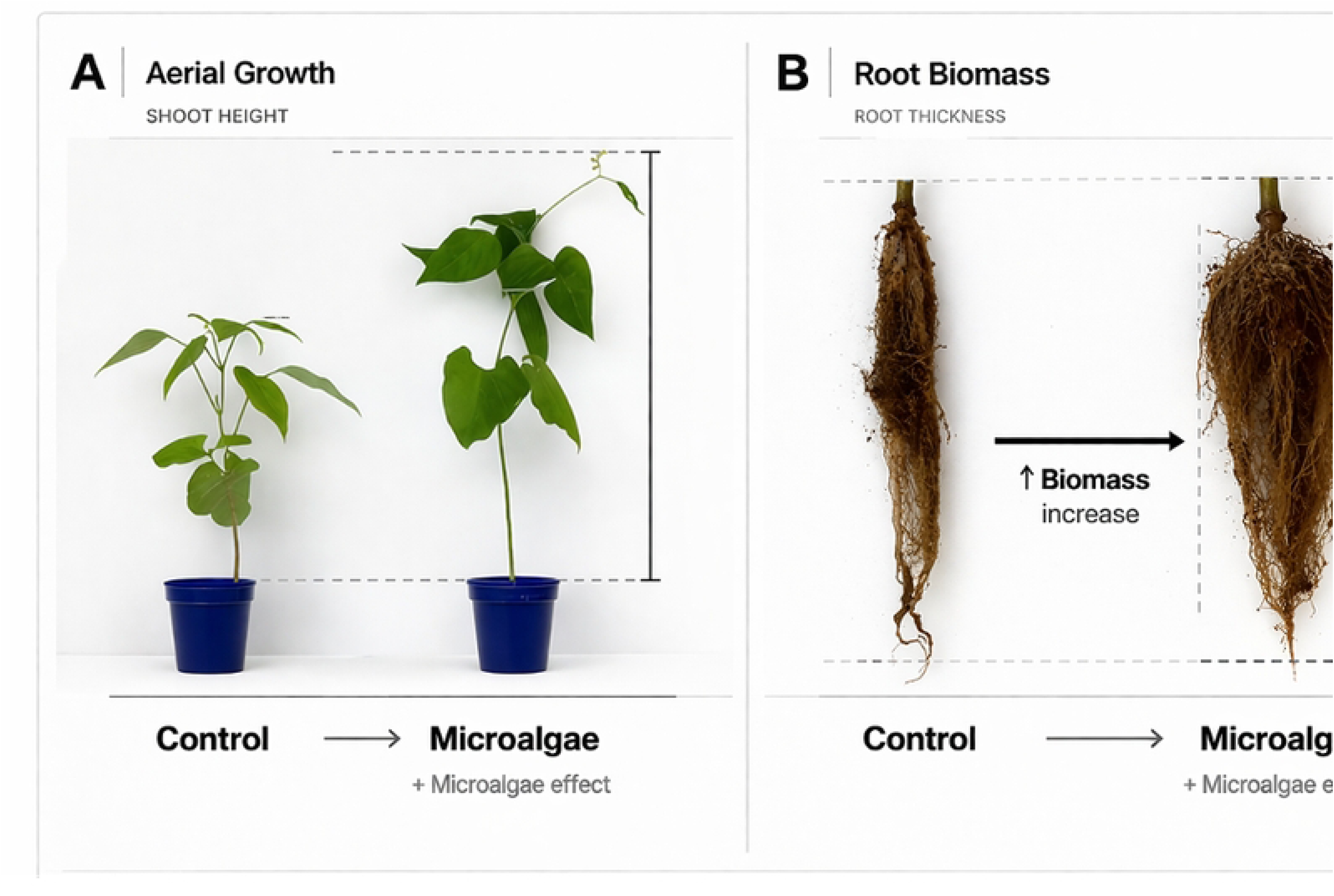
Visible differences in biometric variables due to the application of microalgae. A: height and above-ground biomass; B: root biomass.

### 3.5. Rice

#### 3.5.1. Model performance and optimization

The cubic model provided the best fit for total dry biomass (TDB), explaining 82.3% of the observed variability (Table 3). A significant relationship was found between TDB and the proportions of *C*. *vulgaris*, *Scenedesmus* sp., and *A*. *platensis* (S1 File. Table S3), confirming the suitability of the model for optimization. Response surface analysis identified an optimal formulation composed of 62.3% *A*. *platensis* and 37.7% *C*. *vulgaris*, with no contribution from *Scenedesmus* sp., resulting in a predicted maximum TDB of 1.66 g plant⁻¹ (Fig. 9) (S2 File).

**Fig. 9.**
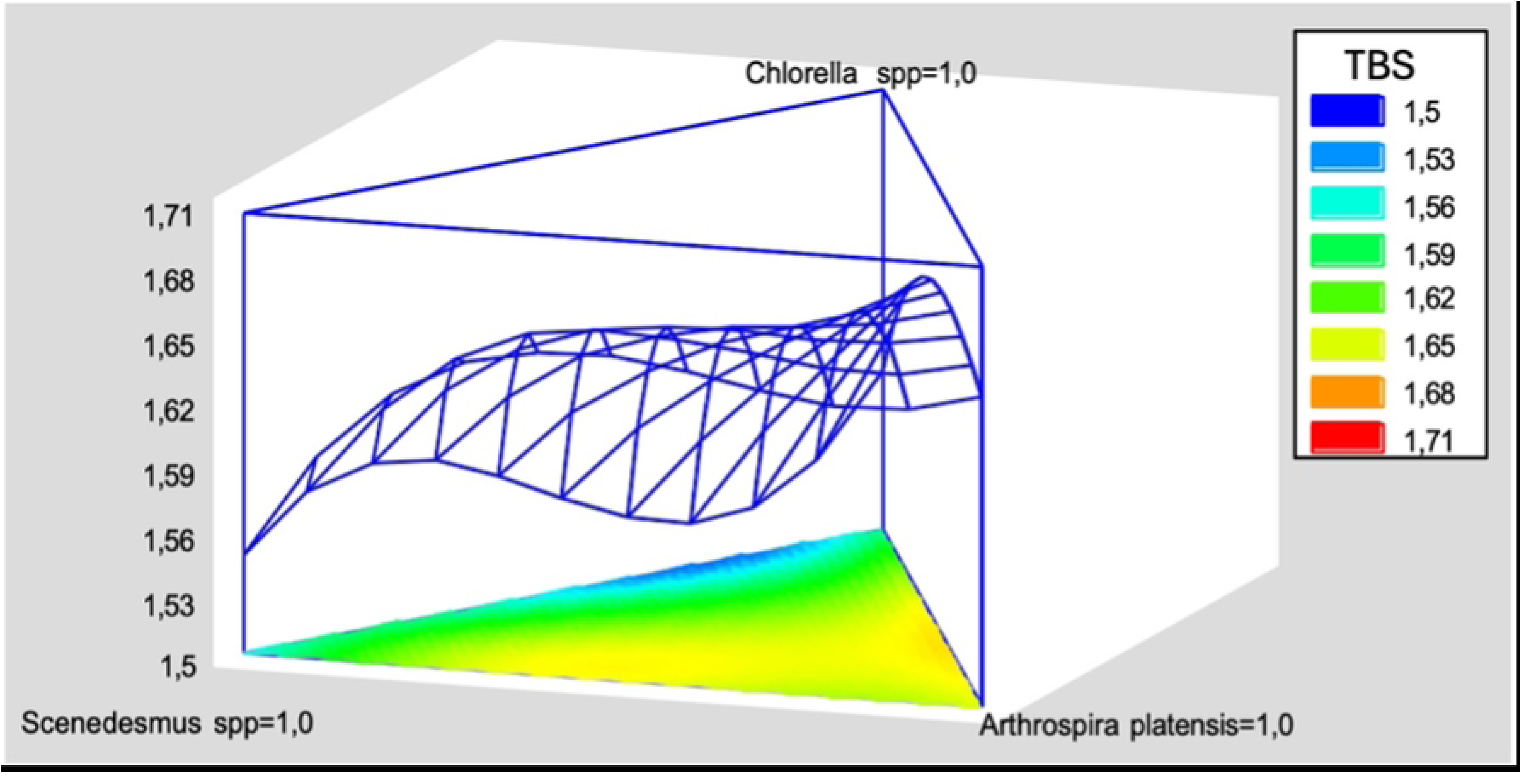
Response surface for TDB indicating the optimal mixture composed of *A*. *platensis* and *C*. *vulgaris*, excluding *Scenedesmus* sp.

**Table 3.** Analysis of variance (ANOVA) for the simplex–lattice mixture model fitted to total dry biomass (TDB).

| *Source* | *Sum*<br>*of*<br>*Squares* | *Df* | *Mean*<br>*Square* | *F-*<br>*Ratio* | *P-*<br>*Value* | *SE* | *R-Squared*<br>*(%)* | *Adj. R-*<br>*Square*<br>*d (%)* |
| --- | --- | --- | --- | --- | --- | --- | --- | --- |
| **Mean** | 73.1272 | 1 | 73.1272 |  |  |  |  |  |
| **Blocks** | 0.00188929 | 1 | 0.00188929 | 0.74 | 0.3962 |  |  |  |
| **Linear** | 0.0320554 | 2 | 0.0160277 | 11.33 | 0.0003 | 0.037610 | 50.00 | **43.75** |
|  |  |  |  |  |  | 5 |  |  |
| **Quadratic** | 0.0148422 | 3 | 0.00494739 | 5.44 | 0.0063 | 0.030163 | 71.86 | **63.82** |
|  |  |  |  |  |  | 8 |  |  |
| **Special** | 0.000274455 | 1 | 0.000274455 | 0.29 | 0.5952 | 0.030685 | 72.26 | **62.55** |
| **Cubic** |  |  |  |  |  | 9 |  |  |
| **Cubic** | 0.00677863 | 3 | 0.00225954 | 3.19 | 0.0310 | 0.026628 | 82.25 | **71.80** |
|  |  |  |  |  |  | 1 |  |  |
| **Error** | 0.0120539 | 17 | 0.000709054 |  |  |  |  |  |
| **Total** | **73.1951** | **28** |  |  |  |  |  |  |

#### 3.5.2. Validation of the optimized formulation

Experimental validation confirmed the predictive performance of the model, as the observed TDB (1.50 g plant⁻¹) closely matched the predicted value. Residual normality and homogeneity of variances were verified for the fitted model (P ≥ 0.05).

Application of the optimized formulation produced a significant biostimulant effect on rainfed rice (P < 0.05), increasing TDB by 12.2% compared with the control (Fig. 10). These differences were also visually reflected in greater shoot and root biomass development of treated plants (Fig. 10 A and B) (S2 File).

**Fig. 10.**
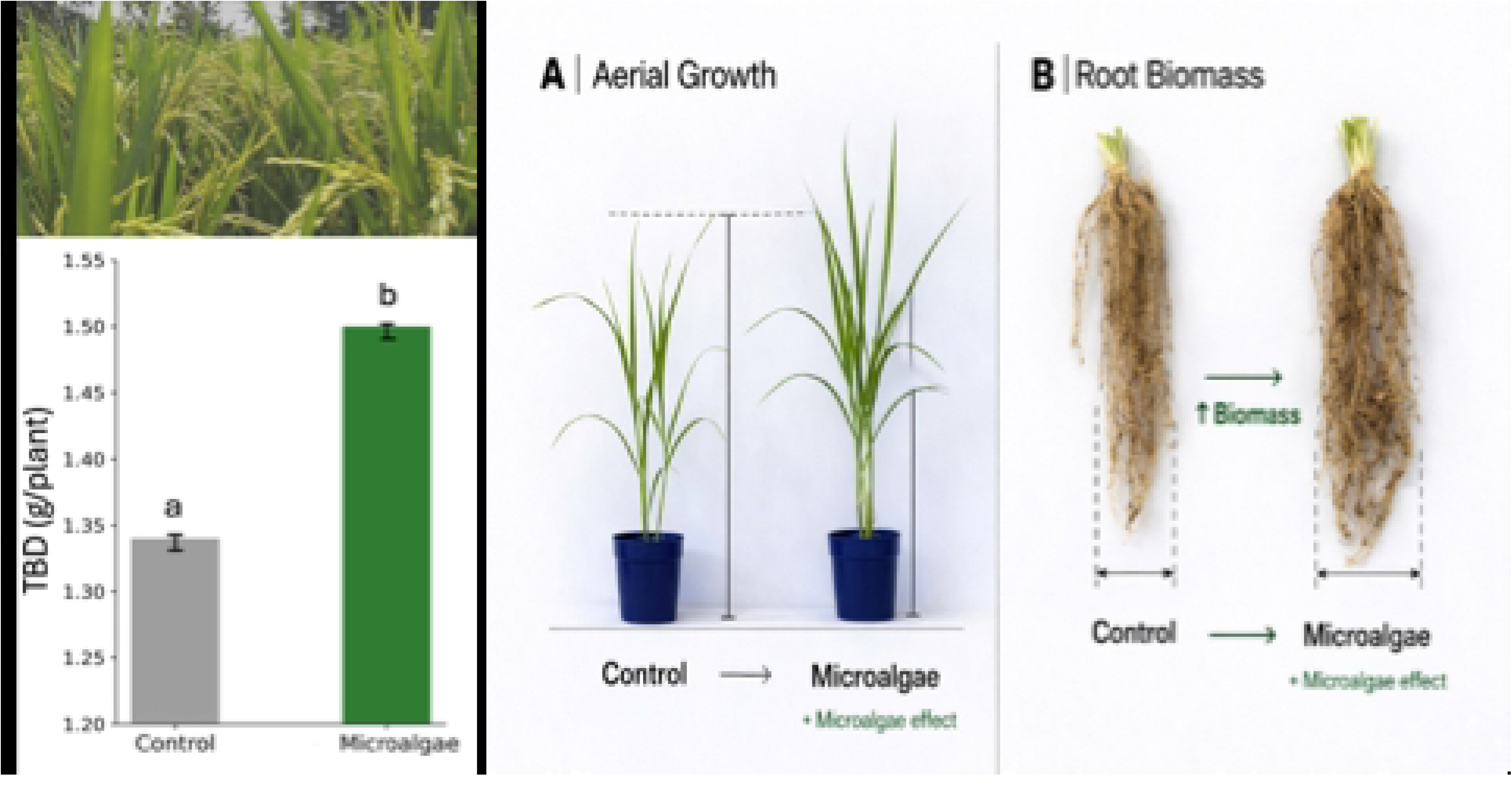
Biostimulant effect of microalgae on TDB. The vertical bars indicate the standard error of the mean (n=10). Means with different letters indicate significant differences according to ANOVA (P ≤ 0.05). Visible differences in biometric variables due to the application of microalgae. A: above-ground biomass; B: root biomass.

## 4. DISCUSSION

The principal contribution of this study is not only the identification of effective microalgal formulations for the two crop species evaluated, but the proposal of an integrated design strategy for the rational development of agricultural formulations based on microalgae and cyanobacteria.

Unlike previous studies, which have predominantly evaluated individual strains or predefined consortia, the proposed strategy integrates biomass production, species characterization, nutritional characterization, mixture design, nutrient profile estimation, biological validation, and statistical optimization into a single workflow. This approach provides a reproducible methodology for designing crop-specific formulations rather than relying on empirical trial-and-error selection.

Although previous studies have demonstrated that microalgae enhance plant growth, nutrient uptake, and soil fertility through multiple mechanisms, including the production of phytohormones and the stimulation of soil microbial communities [13,14], these studies have generally focused on evaluating biological effects rather than establishing an integrated formulation design strategy [15].

A key outcome of the proposed methodology was the identification of distinct optimal formulations for the two crop species evaluated in this study: common bean and rainfed rice, demonstrating that agricultural formulations based on microalgae and cyanobacteria should not be considered universal bioinputs. The formulation optimized for common bean was dominated by *Scenedesmus* sp., whereas the optimal formulation for rice required a higher proportion of *A*. *platensis*. These contrasting formulations indicate that biomass composition should be selected according to crop physiology and desired agronomic responses rather than assuming that a single consortium is suitable for all plant species.

An important feature of the proposed strategy is the incorporation of nutritional characterization during formulation development. Determining the nutritional composition of each biomass allowed estimation of the nutrient profiles of the optimized formulations and their association with crop responses [16]. Although the quantified minerals originated from the culture medium rather than being synthesized by microalgae, their determination provides valuable information for biomass standardization and formulation reproducibility, making nutritional characterization an essential component of formulation development that is generally absent from previous studies.

The differences observed among the optimized formulations are likely explained by the complementary biochemical characteristics of the selected species. Previous studies have shown that *Chlorella* and *Scenedesmus* produce phytohormones such as auxins and cytokinins, together with amino acids and other bioactive compounds that stimulate germination, cell elongation, and root development [17]. Likewise, *Scenedesmus* cultivated in nutrient-enriched media has been reported to increase aboveground and belowground biomass in crops such as pak choi, supporting its potential as an agricultural biofertilizer [18]. In contrast, *A*. *platensis* is characterized by its high nitrogen content (Fig. 3) and the production of polysaccharides, pigments, and other bioactive metabolites that can enhance plant metabolism and stress tolerance. Previous studies have also demonstrated that cyanobacteria release micronutrients [9] and bioactive compounds [19–21] that improve nutrient acquisition and plant performance [22]. These complementary characteristics explain why different proportions of the three species were required to maximize plant performance in bean and rice (Fig. 4).

The crop-dependent responses observed in this study are also consistent with current knowledge regarding plant–microbe interactions within the rhizosphere. Microalgae may directly influence soil microbial communities, which regulate nutrient cycling and bioavailability [23]. Their biomass and extracellular metabolites serve as carbon sources for rhizosphere microorganisms, modifying microbial structure and activity and stimulating processes such as nutrient mineralization and mobilization [24,25]. Consequently, the effectiveness of a given formulation depends not only on the biochemical composition of the microalgal consortium but also on its interaction with the rhizosphere microbiome and the physiological characteristics of the target crop.

Although plants were cultivated under nutrient-sufficient conditions using Hoagland’s nutrient solution, indicating that the observed improvements were primarily associated with biostimulant activity rather than nutrient supply, microalgae are also recognized as potential biofertilizers under nutrient-limited conditions through the gradual release of nutrients and organic matter[16]. Previous studies have shown that *Chlorella sorokiniana* improves plant nutrition and root development in wheat through the combined action of nutrient release and phytohormone production [15].

Similarly, amino acids, polysaccharides, and other bioactive compounds produced by microalgae contribute to improved nutrient use efficiency and increased tolerance to abiotic stress [26]. These observations are consistent with the crop-specific responses obtained in the present study, while reinforcing that formulation composition should be optimized according to the target crop.

From an agronomic perspective, the proposed strategy contributes to the development of next- generation agricultural bioinputs by integrating biomass characterization, formulation optimization, and biological validation into a reproducible design methodology. This concept is consistent with circular bioeconomy approaches that promote nutrient recovery, reduced dependence on synthetic fertilizers, and more sustainable cropping systems [27]. Although validated under greenhouse conditions, additional field studies are required to evaluate formulation performance under different soils, climates, and management systems before large-scale implementation [28].

Overall, this study demonstrates that agricultural formulations based on microalgae and cyanobacteria can be developed through a reproducible design strategy that integrates biomass characterization, mixture optimization, and biological validation. The proposed methodology provides a foundation for the rational development of next-generation microalgal bioinputs and can be adapted to additional crop species in future studies.

## 5. CONCLUSIONS

This study presents an integrated approach for the rational development of agricultural formulations based on microalgae and cyanobacteria by combining biomass production, nutritional characterization, mixture design, nutrient profile estimation, biological validation, and statistical optimization. This workflow provides a reproducible strategy for designing crop-specific formulations rather than relying on empirical selection of individual species or predefined consortia.

The optimized formulations differed between the two model crops, demonstrating that formulation performance is crop-dependent. The optimal formulation for common bean consisted of 31.6% *C*. *vulgaris* and 68.4% *Scenedesmus* sp., whereas rainfed rice required 62.3% *A*. *platensis* and 37.7% *C*. *vulgaris*. These findings highlight the importance of crop-specific formulation design for the two crop species evaluated in this study and provide a methodological basis for developing next- generation agricultural bioinputs based on microalgae and cyanobacteria.

## Acknowledgments

The authors gratefully acknowledge the Universidad de Antioquia and Minciencias for their support of this research.

## Supporting information

S1 File. Supplementary data tables.

S2 File. The equations of the fitted cubic models for each variable.

## Authors contributions

Carlos Lopera: Methodology. Madelem Giraldo: Methodology, Investigation, Formal analysis and Conceptualization. Natalia Herrera: Methodology, Investigation, Formal analysis and Conceptualization. All authors read and approved the manuscript.

